# Analyzing Biodiversity in Protected vs. Unprotected Tide Pools

**DOI:** 10.1101/2024.10.03.616586

**Authors:** Vicki Chen, Rowan Campbell, Arohi Chirputkar, Emma Tran, Parnika Chaturvedi, Sally Han, Amber Lu, Zoe Chu, Melody Erb, Tejin Mehta, Vidya Bindal, Isaac Lee, Andrew Benson, Connor T. Adams

## Abstract

This study assessed the efficacy of marine protected areas (MPAs) by comparing tide pool biodiversity in Half Moon Bay, California between a protected site (Fitzgerald Marine Reserve) and an unprotected site (Maverick’s Beach). Samples were collected through random quadrat surveys. ImageJ and RStudio were used to analyze data through three methods utilizing the Simpson’s Diversity Index along with three metrics. The unprotected Maverick’s Beach displayed greater biodiversity through two metrics while the third did not find significant difference in biodiversity between sites. 30 unique species were present at each site with 43 species between both. These mixed findings prevent us from drawing definitive conclusions on the effectiveness of Half Moon Bay’s tide pool MPAs but suggest the MPA is less biodiverse. Potential explanations include the Intermediate Disturbance Hypothesis, lack of sample size, or that the MPA is not effective in maintaining biodiversity.

## Introduction

To evaluate the potential fragility of marine ecosystems exposed to human activities, the need for effective methods to monitor species richness and diversity in these environments has become essential. One key approach to maintaining healthy ecosystems has been the designation of marine protected areas (MPAs), which are designed to prohibit anthropogenic activities in order to provide ecosystems time to recover or create a core region for species recruitment. The human impact on tide pools remains to be fully elaborated, but negative outcomes could be possible through, e.g., the trampling of habitats and the harvesting of local organisms (Werfhorst and Pearse 2007). To counteract potential negative impacts of human activities in sensitive ecosystems, California, like many other regions with rich coastal ecosystems, has implemented MPAs which are defined as a “discrete geographic marine or estuarine area seaward of the mean high tide line or the mouth of a coastal river, including any area of intertidal or subtidal terrain, together with its overlying water and associated flora and fauna that has been designated … to protect or conserve marine life and habitat. An MPA includes marine life reserves and other areas that allow for specified commercial and recreational activities, including fishing for certain species but not others, and kelp harvesting… MPAs are primarily intended to protect or conserve marine life and habitat, and are therefore a subset of marine managed areas (MMAs), which are broader groups of named, discrete geographic areas along the coast that protect, conserve, or otherwise manage a variety of resources and uses, including living marine resources, cultural and historical resources, and recreational opportunities.” (Marine Life Protection Act - FGC § 2852). Classifications of MPAs include marine life reserves (the equivalent of the state marine reserve classification), state marine parks (which allow recreational, but not commercial fishing), and state marine conservation areas (which allow for specified commercial and recreational activities, including fishing for certain species and kelp harvesting, provided that these activities are consistent with guidelines of the area). Positive side effects of MPAs may include the dispersal of fish larvae and adults outside the boundary of mentioned MPAs, which can benefit nearby fisheries and other non-MPA zones by providing areas of strong recruitment that increases catch along MPA borders as individuals migrate out. This increase in abundance has been shown to increase fishery productivity despite the decrease in fishable areas (Harrison et al. 2012). MPAs that prohibit commercial activity and overexploitation in the habitat are designated as “no-take MPAs” (Bennett 2014). While no-take MPAs have been shown to be the most effective, some MPAs will permit light fishing (Sala, Giakoumi 2018).

It may also be useful to examine how human impact differs between intertidal zones. The intertidal zone is the area of land exposed at low tide which is covered at high tide. The high intertidal zone refers to land that is dry for most of the day except during high tide. The low intertidal is submerged except during spring tide when the tide is at its lowest. The mid intertidal zone is the land between these two.

Quantitative methods for biodiversity and species community structure are needed to monitor if MPAs are effective in protecting ecosystems. Biodiversity includes several factors: species richness (the number of unique species present) and relative abundance (the number of individuals of each species present), which are measured to calculate species evenness. Simpson’s diversity index is a value that accounts for both richness and relative abundance to quantify the diversity of a community. The equation for acquiring this value is as follows:

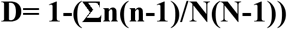

D: Diversity Index

n: number of individuals of a single species

N: number of individuals in a population

The values of Simpson’s Diversity Index range from 0 to 1, with higher values indicating greater biodiversity. In theory, effective MPAs will have more species diversity, larger average size for many species, and higher populations when compared to an unprotected area. For example, a study on the efficacy of MPAs within the Northern Channel Islands reserve network showed that species biomass within California’s MPAs increased at higher rates than outside the MPAs (Murray and Hee 2019). However, this question is not fully resolved across different types of MPAs and variation in MPA impacts might be expected across regions or contexts. In the current study, we hypothesized that higher human activity in non-protected tide pool areas would be associated with reduced biodiversity, average size, and abundance of local species when compared to an MPA tide pool.

## Materials and Methods

### Study Site Information

We visited the MPA tide pool Fitzgerald Marine Reserve, Moss Beach, California (37.49571178714731, -122.49798770000001; latitude and longitude) on 2 occasions (7/15/2021, 7/27/2021), and the unprotected tide pool at Mavericks Beach, Half Moon Bay, California (37.52403303453021, -122.51758327301393) on 3 occasions (10/20/2021, 12/4/2021, 2/27/2022) to conduct random sampling on species counts, sizes, abundance, and biodiversity quantifications in the area. Both are located in San Mateo County, CA, 4.4 km apart. By choosing sites in close proximity it is hoped that factors other than regulatory status can be controlled for. The high, mid, and low intertidal zones were sampled to collect a variety of data, and upwards of 2-3 hours was spent at each location.

### Quadrats

A series of quadrats were placed, and the species within were then identified and recorded. We used passive quadrat sampling, meaning the organisms found within the quadrat were not removed. At each site visit, photographs were taken for later analysis through ImageJ (described below). To place the quadrats, two transect lines were set, which served as the x and y axis. The x-axis was 50 meters North/South, and the y-axis was 30 meters East/West. A random number generator established several coordinates along these axes to indiscriminately select quadrat sampling locations. New Quadrats were selected during each visit, rather than sampling the same quadrats each visit. 37 unique quadrats were sampled at Fitzgerald Marine Reserve and 57 quadrats were sampled at Mavericks Beach. Additionally, five targeted locations were surveyed at Maverick’s Beach due to unique species and high biodiversity. These targeted quadrats included 1 area where sea stars were abundant, 3 densely populated tide pools, and 1 quadrat on rocky substrate. These 5 quadrats are included in the public dataset but were not used during analysis due to their selection being nonrandom and only at one location. In the dataset they are labelled as targeted locations (Adams, Benson 2024). 50×50 cm quadrats were placed at the selected locations. Several photos were taken at each quadrat, starting with a picture of the entire quadrat, and then zooming in on individual portions for more detailed and accurate identification. To get a better view of quadrats that were overgrown with plant life, algae and seagrass were moved away from the frame after recording data with the mentioned vegetation. A ruler was included in the frame for each picture to establish scale. The camera was positioned at the same angle above the quadrat to ensure the scale would not be warped. The sampling technique did not allow for complete coverage of specific areas such as mussel beds, as they were inaccessible to the lines.

### Analyzing the Quadrats

Quadrat images were analyzed using ImageJ image processing software developed by the National Institute of Health. The close-up pictures of each quadrat allowed for identification of the species present. The size of each organism was measured with ImageJ by setting a scale using the ruler placed in each quadrat photo. Specifically, the size of organisms (excluding those that could not be discreetly counted, i.e seagrass and algae) was considered to be the longest length across the organism in cm. To measure the percentage cover of organisms such as black turban snails and mussels, the freehand selection tool or oval selection tool was used to determine their area in cm^2^ and divided the value by the area of the quadrat, 2500 cm^2^. To quantify the percentage cover for algae, seagrass, or other species that cannot be discreetly counted, a picture that showed a total view of the entire area was used; with the grid nature of the quadrats, we were able to get an estimate of percentage cover given that each grid represents 4% of the total quadrat area.

Data were inputted to a master spreadsheet that included the area visited, coordinates, intertidal zone, as well as the name and number of each species with their corresponding lengths. RStudio was used for statistical analysis and graph creation. Data was grouped by quadrat and sums found for species within quadrats. This was used to generate lists that the simpson function in RStudio’s abdiv package could use to find the Simpson diversity value for sites, tidal zones, and individual quadrats as data was grouped by these parameters. Statistical analysis on the differences in biodiversity index values between sites and intertidal zones was done by finding the Simpson’s Index value for each quadrat, separating the list of results by site and tidal zone, then using RStudio’s t.test function comparing corresponding lists. Finally, we tested for differences between the percent coverage of all organisms. A copy of the datasheet was edited to include only 1 instance of a species with an associated percent cover value equal to the sum of all instances of that species within the quadrat. The percent cover value of each species in the quadrat was used with RStudio’s simpson function to find a diversity value for each quadrat, and those values were used with the t.test function to compare biodiversity by percent coverage between sites and intertidal zones.

The only species numerous enough at both sites to analyze difference in average length was the Black Turban Snail (*Tegula funebralis*). RStudio’s t.test function was used with lists of Black Turban Snail lengths from each site.

This study used three methods to determine biodiversity using the Simpson’s Diversity Index. Method 1 omitted all species that could not be discreetly counted as individual organisms from the species richness count. Examples included seagrass and algae. Method 2 counted each instance of an organism unable to be counted discreetly in a quadrat as one. For example, *Saccharina sessilis* visible in a quadrat would be counted as one instance of the organism regardless of percent coverage. Method 3 considers only the percent coverage of each species. However, due to technical issues method 3 could not be applied to all quadrats, and so was performed with a smaller sample size. 53 quadrats at Maverick’s Beach and 24 quadrats at Fitzgerald Marine Reserve.

## Results

In all quadrats, there were a combined 43 unique species. 30 at Mavericks Beach and 30 at Fitzgerald Marine Reserve.

**Table 1:**
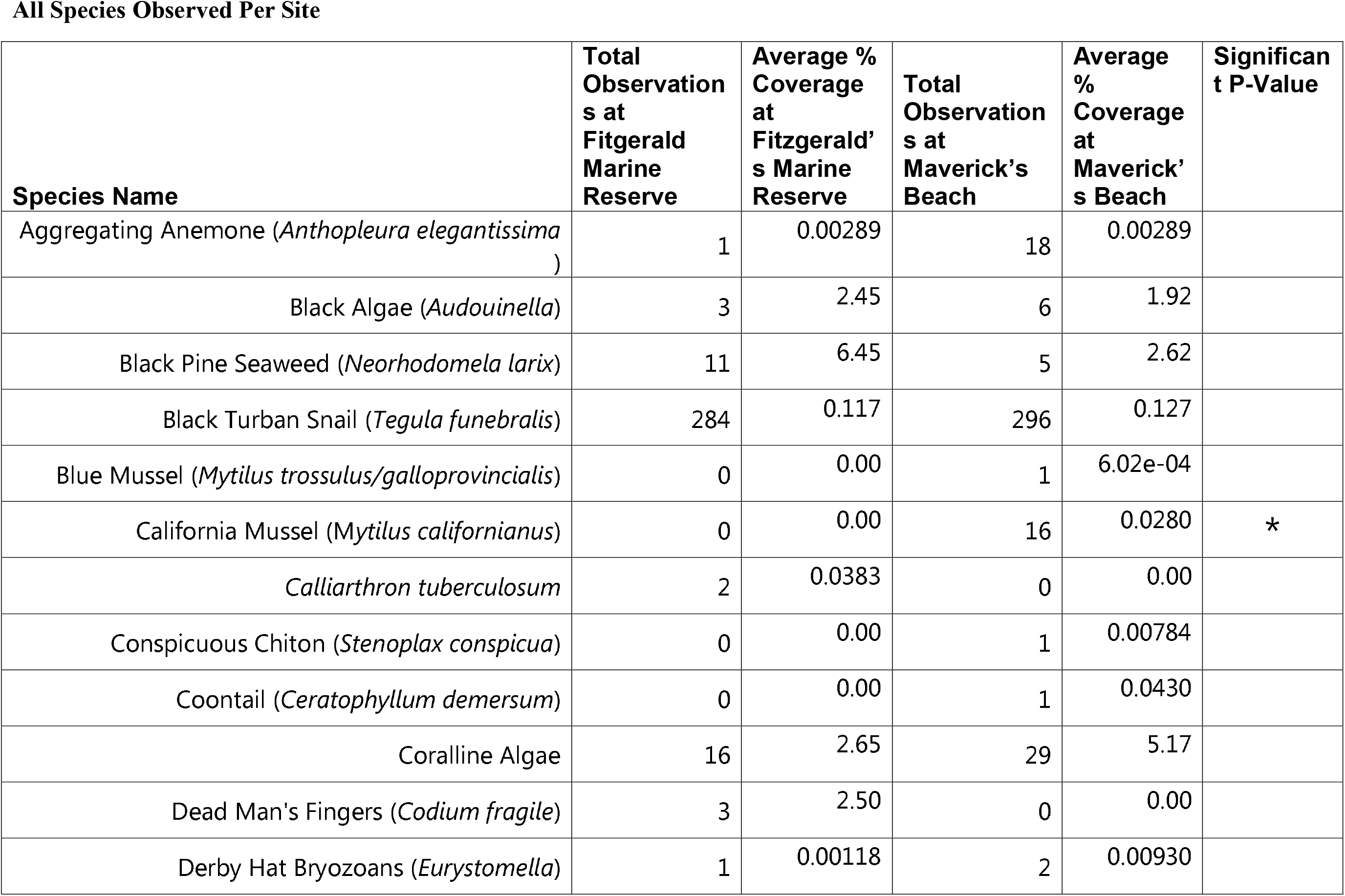
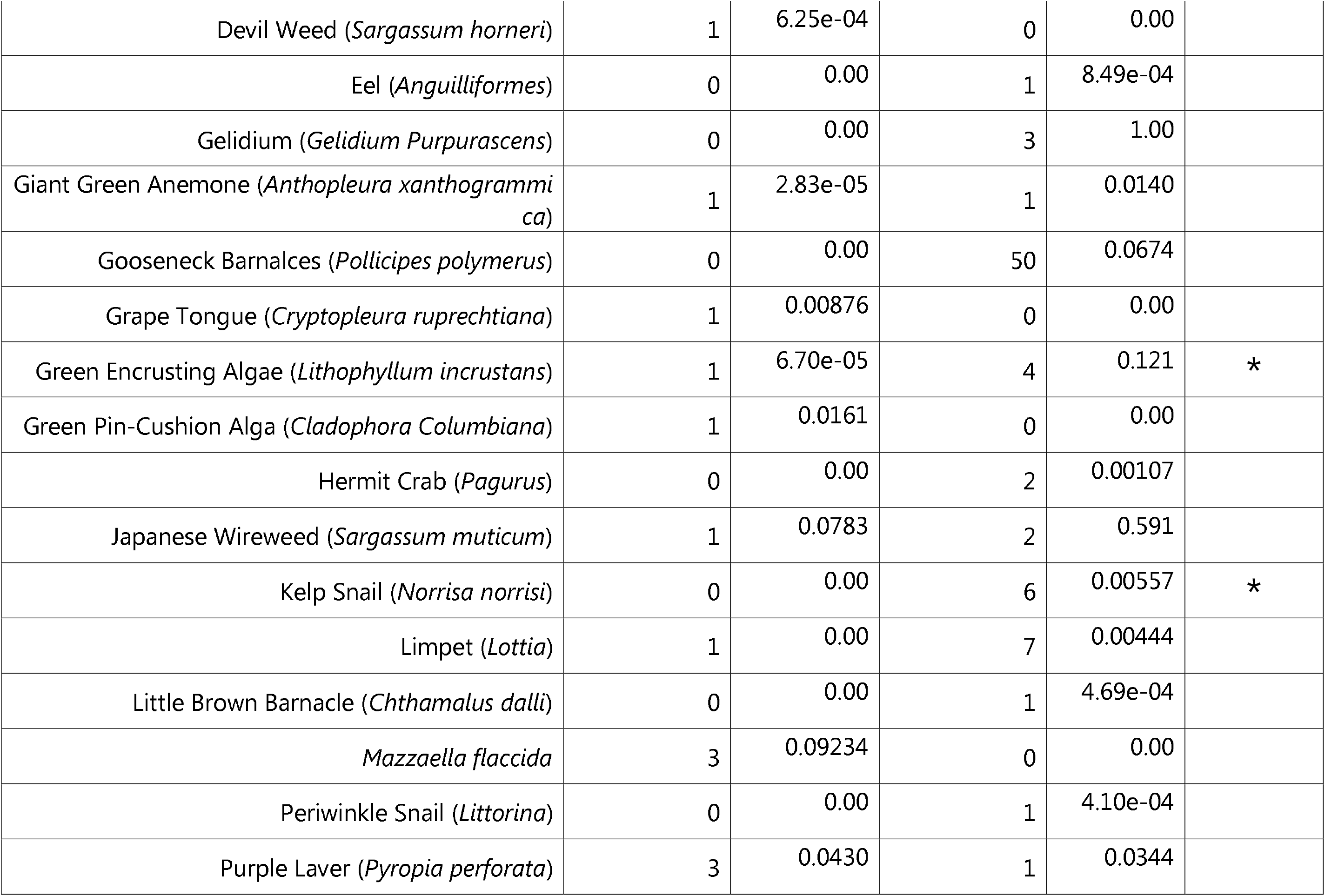
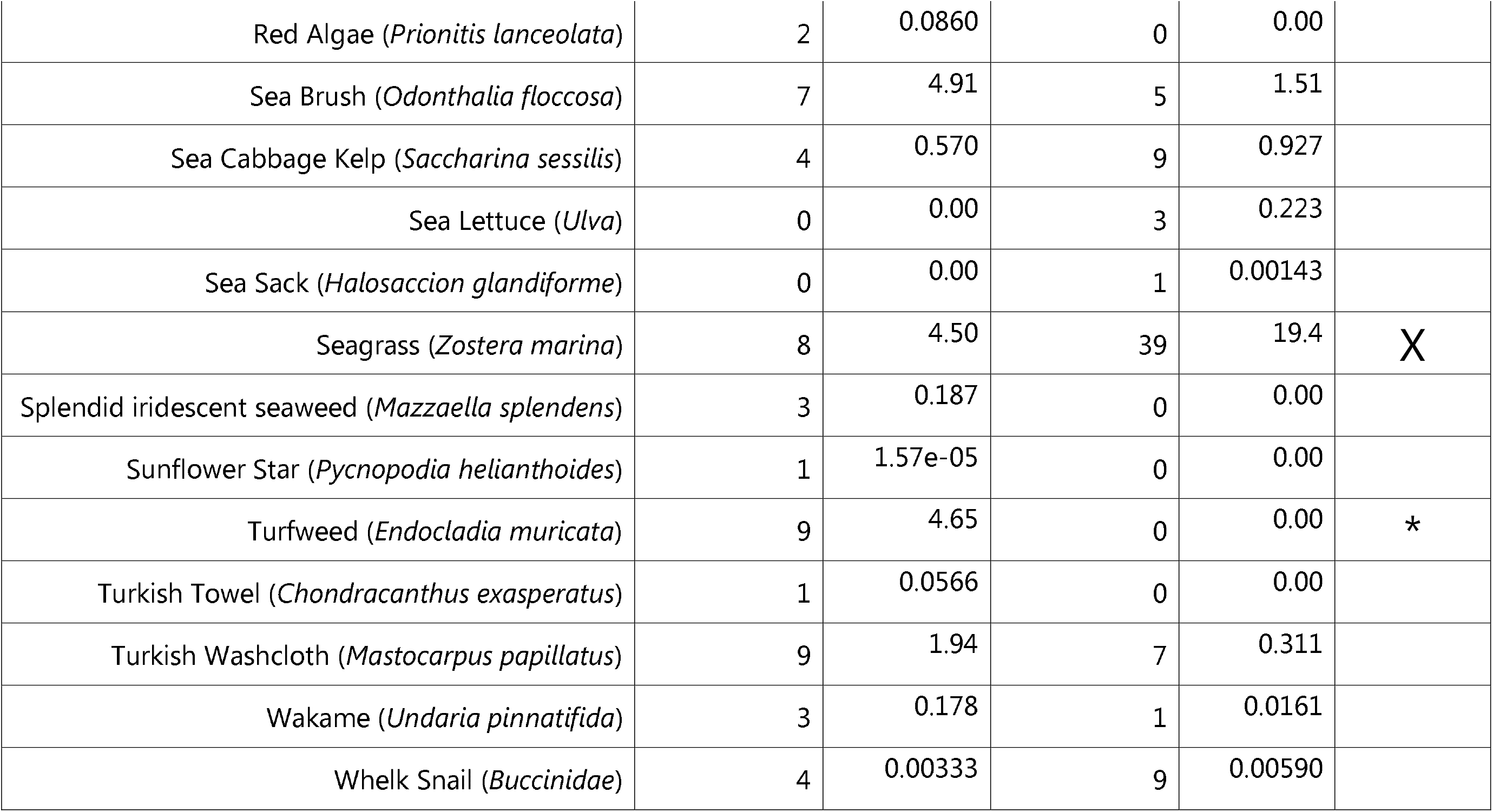
This table contains a row for each species observed at either site. Column 1 contains species names. Columns 2 and 4 show how many individuals were observed at Fitzgerald Marine Reserve and Maverick’s Beach respectively. Species which could not be counted as individuals depict how many quadrats the species were in instead. Columns 3 and 5 depict the average % coverage of the species at each site. Column 6 notes if the difference in % coverage between sites was significantly different. * Represents p<0.05. X Represents p<0.001.

Among those species observed, most did not feature statistically significant differences due to either similarity or small sample size. Running t-tests on the significance of size and percent coverage for each individual species yielded mostly insignificant results. The following featured statistically significant differences in percent coverage.

Seagrass (*Zostera marina*) coverage between the sites was found to be significant p=7.532e-5. Mean coverage was 29.93% at Maverick vs 7.862% at Fitzgerald.

Turfweed (*Endocladia muricata*) was significant: p=0.01265. *Endocladia muricata* was only present at Fitzgerald. Average coverage at Fitzgerald 4.588%.

Green Encrusting Algae (*Lithopyllum incrustans*) was slightly significant p=0.04635. Average coverage was 3.005e-1% at Mavericks vs 6.627e-5% at Fitzgerald.

California Mussel (*mytilus californianus*) was only found in Mavericks. p=.02662 average percent cover at Mavericks .05647%

Kelp Snail (*Norrisa norrisi*) was only present at Mavericks. p=0.02864 average percent cover at Mavericks 0.007319%.

Species richness per quadrat was compared between sites and shown to not be significantly different, p=.07562. Comparing intertidal zones was also insignificant.

For all three methods of measuring the Simpson’s diversity index, the Simpson value was found for each site and its tidal zone overall, as well as for each quadrat within those sites. The index value of individual quadrats was used for comparative statistical analysis between sites.

Method 1, where species that cannot be counted are dropped from the calculation, yielded a higher average diversity value per quadrat at Mavericks (p=0.009911). The differences between the high (p=.2546) and low (p=.3142) intertidal zones were not statistically significant, but the mid intertidal zone was significantly different p=.01408. For quadrats in the mid intertidal zone, Mavericks had an average Simpson’s index value of .2441 while Fitzgerald had .08014.

**Table 2.** Simpson’s Diversity Index values are distinguished by location and tidal zone for Method 1. Overall values are listed for intertidal zones.

|  | Mavericks | Fitzgerald |
| --- | --- | --- |
| Overall | 0.6171799 | 0.1276698 |
| Average Per Quadrat | 0.21806179 | 0.08024407 |
| Low Intertidal | 0.4310532 | 0.1276698 |
| Mid-Intertidal | 0.712707 | 0.1762695 |
| High Intertidal | 0.2590587 | 0 |

Method 2, where species that cannot be discreetly counted are given a value of 1 in each quadrat in which they appear, did not yield a significant difference in quadrat diversity between sites (p=0.6084). Similarly, none of the intertidal zones displayed a significant difference between each other with a p-value of 0.3245 in the low intertidal zone, 0.1277 in the mid intertidal, and .4496 in the high intertidal.

**Table 3.**
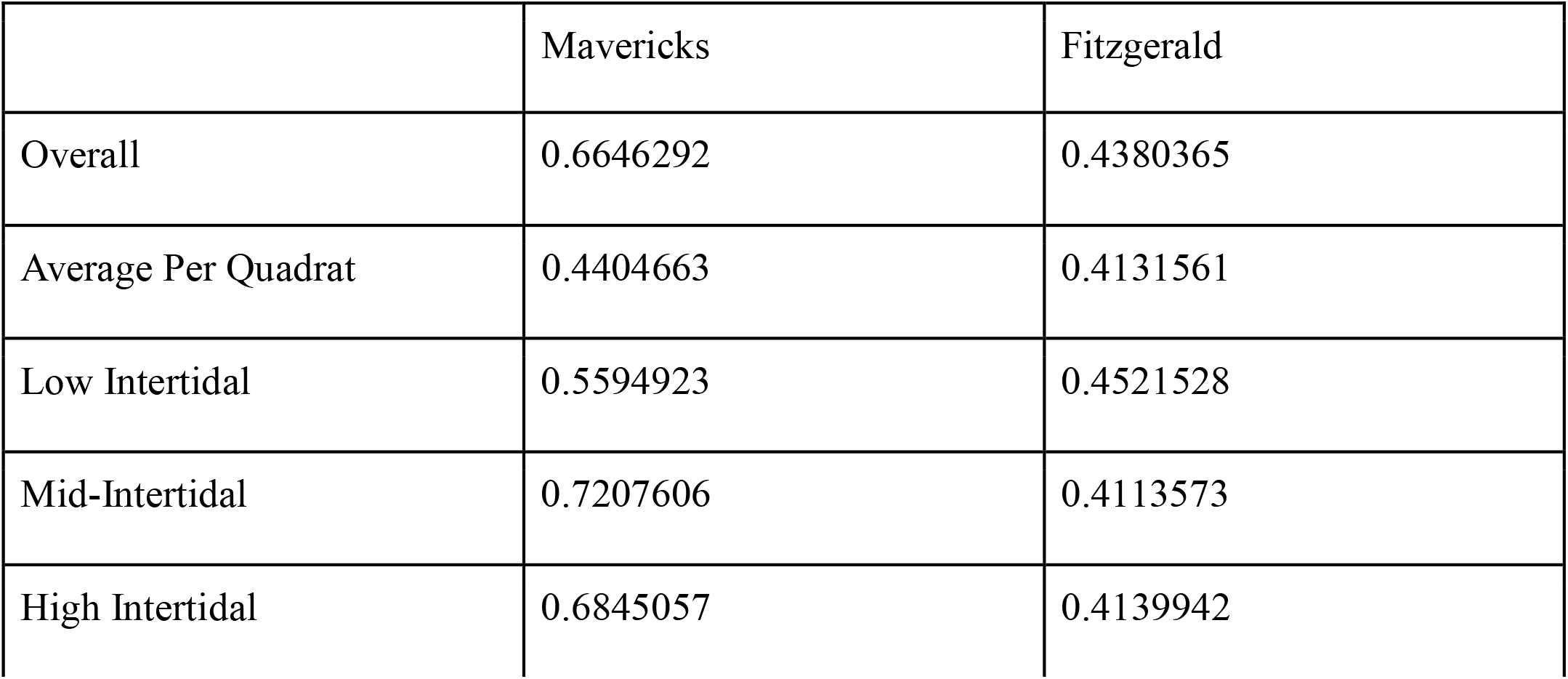
Simpson’s Diversity Index values are distinguished by location and tidal zone for Method 2. Overall values are listed for intertidal zones.

Method 3, where percent cover was used to calculate biodiversity rather than individual count, yielded a higher average biodiversity value per quadrat at Mavericks p=0.0357. Despite having a lower average Simpson’s value per quadrat, Fitzgerald had a higher index score overall. Because statistical analysis can only be performed on the per quadrat values, they were given more credence in our conclusions. The difference in index values between intertidal zones was not significant.

**Table 4.** Simpson’s Diversity Index values are distinguished by location and tidal zone for Method 3. Overall values are listed for intertidal zones.

|  | Mavericks | Fitzgerald |
| --- | --- | --- |
| Overall | 0.6994125 | 0.8732088 |
| Average Per Quadrat | 0.2927487 | 0.1886056 |
| Low Intertidal | 0.601344 | 0.8573748 |
| Mid-Intertidal | 0.7039219 | 0.7701544 |
| High Intertidal | 0.5089866 | 0.7279662 |

There was a significant difference between the size, but not the number or percent coverage, of black turban snails between sites. A t test on the difference in length found black turban snails at Mavericks beach to be larger p=1.218e-12. Means sizes are 1.389 cm at Mavericks vs 1.050 cm at Fitzgerald

**Figure 1:**
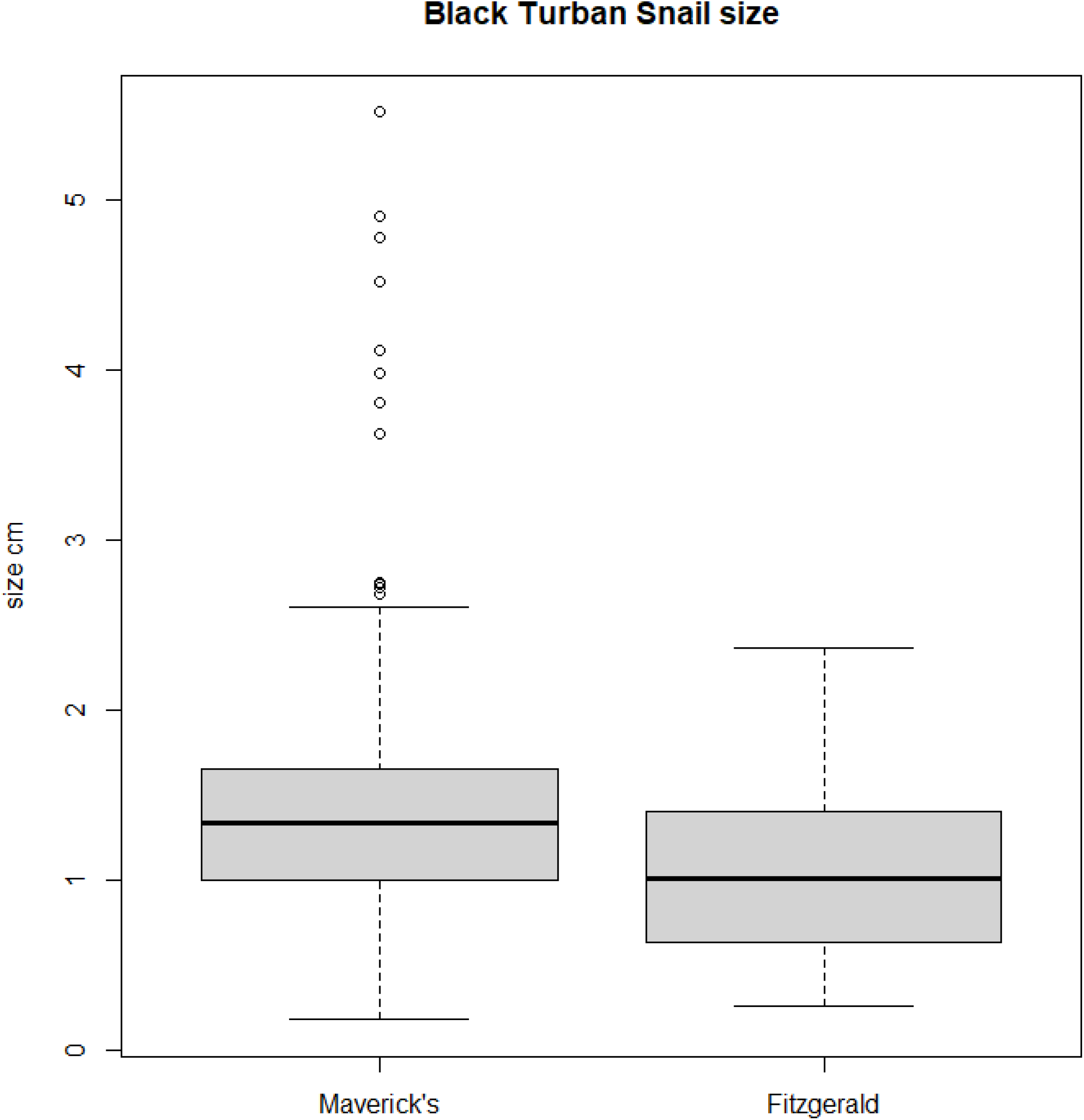
Boxplot comparing black turban snail sizes between Maverick’s Beach and Fitzgerald Marine Reserve

## Conclusion

Our study does not confirm that Half Moon Bay’s tide pool MPAs are quantifiably more effective at promoting biodiversity when compared to unprotected tide pools. The results show that the tide pools in the MPA, Fitzgerald’s Marine Reserve, had a significantly lower biodiversity rating than that of the non-MPA through measurement methods 1 and 3, and were not significantly higher through method 2. Equal numbers of unique species were observed at both sites, and there was no significant difference in average species richness per quadrat between sites. This was unexpected, as the goal of MPAs is to preserve biodiversity, with previous studies reporting success in this regard. Potential explanations for these results may include the MPA being ineffective or poorly enforced, the unprotected area not being greatly affected by human visitors, or the results may support the Intermediate Disturbance Hypothesis.

The intermediate disturbance hypothesis states that a community with little to no interference (such as natural disasters, human impact, and introduction of invasive species) will develop minimal species diversity because the most advantageous species within a niche will outcompete and exclude others within their niche. In contrast, a community with high disturbance rates will exclude those species unsuited to harsh conditions created by disturbances, in favor of species able to either weather the disturbance or colonize the environment quickly after. However, if a community is subject to occasional intermediate-level disturbances, species diversity is expected to be at its peak because all organisms will experience some disturbance, preventing any one species from remaining dominant long enough to exclude its competitors completely (Connell, 1978). Our team hypothesizes that the Intermediate Disturbance Hypothesis is a potential explanation for why the results did not align with our hypothesis, and the unprotected tide pool featured greater biodiversity than the protected MPA through two of three metrics. The Simpson’s diversity index value may be higher at Maverick’s tide pools because its community is closer to intermediate-level disturbance than that at Fitzgerald Marine Reserve, causing some organisms to be excluded from the site. We hypothesize that because there are heavy regulations on fishing and overall human presence, the community at Fitzgerald experiences little to no disruption, allowing for certain species to thrive and dominate the community.

Alternatively, the fact that Mavericks Beach is not significantly more diverse by all metrics may indicate that the difference in biodiversity between sites is not large. Biodiversity was closest when uncountable species were treated as one instance. This may imply that biodiversity is similar, but the unprotected site is covered by a smaller variety of algae and coralline species. This could be due to the dominant cover species at Mavericks being those more resistant to human trampling.

We observed greater Seagrass (*Zostera marina)* coverage at the unprotected tide pool, likely because seagrass is well adapted as a colonizing species due to its rapid growth rate and resistance to trampling.

Though the population size of Black Turban Snails (*Tegula funebralis*) measured by both number of individuals and percent cover was not significantly different between the sites, average snail size was, with the unprotected site featuring greater average snail size. This may be due to the dominance of Seagrass (*Zostera marina*) and at the site, as it is a potential food source for the herbivorous Black Turban Snails that consume macro algae and organic detritus as primary food sources (Best, 1964). The greater abundance of Seagrass to eat at the unprotected site may allow the black turban snails to consume more and grow larger than those at the MPA. Another possible explanation is that black turban snails are not commonly harvested for consumption, and so may thrive when potential competitors are removed by fishing and human take. Black turban snails may also be resistant to trampling compared to other soft bodied organisms.

Comparing intertidal zones, only the mid intertidal zone in method one featured a significant difference. This may be due to the small sample size once each site was split by zone. That the mid intertidal zone is the only site to feature significant differences even when sites overall were significantly difference may suggest the mid intertidal zone is the most affected by human impact. Our team hypothesizes this may be because the low intertidal zone is most often submerged beyond visitor reach, while the high intertidal zone does not attract as many visitors who are more drawn to the tide pools closer to the ocean which by most of our measurements had greater biodiversity.

## Supporting information

https://data.mendeley.com/datasets/gnjkj3bhg5/1

## Funding Sources

This project was funded by the Aspiring Scholars Directed Research Program (ASDRP). ASDRP had no impact upon the design, collection, analysis, or interpretation of the data.

### Co-Author Contributions

- Andrew Benson: Conceptualization, data curation, project administration
- Connor Adams: Formal analysis, project administration, investigation, visualization, writing
- Vicki Chen, Rowan Campbell, Arohi Chirputar, Emma Tran, Issac Lee, Ian Chen, Parnika Chaturvedi, Akilan Dorairaj, Zoe Chu, Melody Erb, Tejin Mehta, Vidya Bindal: Data curation, analysis, investigation

